# Rapid conversion of replicating and integrating *Saccharomyces cerevisiae* plasmid vectors via Cre recombinase

**DOI:** 10.1101/2020.11.03.367219

**Authors:** Daniel P. Nickerson, Monique A. Quinn, Joshua M. Milnes

**Affiliations:** Department of Biology, California State University, San Bernardino, CA 92407; Department of Biochemistry, University of Washington School of Medicine, Seattle, WA 98195-3750; Washington State Department of Agriculture – Plant Protection Division, Yakima, WA 98902

**Keywords:** Cre recombinase, homologous recombination, plasmid, shuttle vector, overexpression, molecular cloning, genetic engineering, yeast

## Abstract

Plasmid shuttle vectors capable of replication in both *Saccharomyces cerevisiae* and *Escherichia coli* and optimized for controlled modification *in vitro* and *in vivo* are a key resource supporting yeast as a premier system for genetics research and synthetic biology. We have engineered a series of yeast shuttle vectors optimized for efficient insertion, removal and substitution of plasmid yeast replication loci, allowing generation of a complete set of integrating, low copy and high copy plasmids via predictable operations as an alternative to traditional subcloning. We demonstrate the utility of this system through modification of replication loci via Cre recombinase, both *in vitro* and *in vivo*, and restriction endonuclease treatments.

## INTRODUCTION

Yeast (*S. cerevisiae*) has long served as a premier experimental system for exploring eukaryotic genetics and cell biology (Botstein and Fink 2011) and has more recently emerged as a preferred system in synthetic biology (Lee *et al.* 2015). A key component of the yeast genetic toolkit is the availability of shuttle vectors, plasmids capable of replication and selection when introduced into either a yeast or *E. coli* cell (Sikorski and Hieter 1989; Da Silva and Srikrishnan 2012). In their original design, yeast shuttle vectors would typically be modified *in vitro* and subsequently propagated in bacterial cells to generate a desired gene construct that could be later introduced into yeast to study eukaryotic cell biology. In current practice the classical bacteria-then-yeast workflow is often turned on its head by gene synthesis techniques that capitalize upon the readiness of yeast to piece together compatible DNA fragments via homologous recombination (Gibson 2009), so initial gene synthesis and molecular cloning steps often start inside the yeast cell.

Yeast shuttle vectors vary in their selectable markers, typically genes for auxotrophic rescue or antibiotic resistance, and in the presence or absence of a yeast replication locus. Yeast integrating plasmids lack an independent replication locus and must be incorporated into a chromosome in order to replicate. Low copy, centromeric plasmids contain an autonomous replicating sequence (ARS) to initiate DNA replication and a centromere (*CEN*) to support plasmid inheritance during cell division. High copy, episomal vectors carry the *2μ* circle replicating origin. Low and high copy plasmids offer convenience for gene expression studies and functional analyses, but both varieties of replicating plasmids have demonstrated instability in their copy numbers (Futcher and Carbon 1986; Mead *et al.* 1986; Resnick *et al.* 1990). When considering the possible toxicity of plasmid-encoded gene or combination of genes, or even the energetic cost of maintaining the plasmid itself (Mead *et al.* 1986), unstable copy numbers of centromeric and 2μ plasmids can present clear obstacles to experimental execution and interpretation. To ensure even copy number and consistent level of gene expression, some plasmid-encoded genes must be integrated into a chromosome, but such necessity is often realized after experimentation with expression from replicating plasmids. Ability to switch between low copy, high copy and integrating shuttle vector strategies is a key feature of the yeast genetics toolkit.

Previous efforts in improving resources for yeast shuttle vectors have focused heavily on engineering sequences flanking the yeast selectable marker loci to offer convenient removal and remodeling (Gueldener *et al.* 2002; Carter and Delneri 2010; Fang *et al.* 2011; Chee and Haase 2012; Agaphonov and Alexandrov 2014; Siddiqui *et al.* 2014; Jensen *et al.* 2014), either in the plasmid or after chromosomal integration. By comparison, the replication loci of the shuttle vectors have been neglected. We believe a similar level of utility is achieved by engineering the yeast replication loci of shuttle vectors to be removed and remodeled in analogous manner.

We have modified the popular family of pRS shuttle vectors (Sikorski and Hieter 1989; Christianson *et al.* 1992) plasmid shuttle vectors to include yeast replication loci flanked by LoxP sites or triplicate endonuclease cut sites. We demonstrate the utility and flexibility of this system through in vivo and in vitro remodeling of the yeast replication loci and the rapid generation of a complete suite of integrating, low copy and high copy plasmids to support functional analysis in yeast.

## RESULTS & DISCUSSION

### pDN5xx & pDN6xx vector series

While several families of yeast shuttle vectors have been deployed over nearly four decades (reviewed in Da Silva and Srikrishnan 2012), availability of the pRS series of vectors (Sikorski and Hieter 1989; Christianson *et al.* 1992) was a landmark in yeast genetics; the plasmids were rapidly adopted by the field and their use remains ubiquitous due to several useful features (Figure 1 A): i/ yeast auxotrophic selection markers (*HIS3, TRP1, LEU2* or *URA3*) are compatible with engineered auxotrophies in many of the most commonly used yeast laboratory strains (Sherman 2002); ii/ plasmid selection in bacteria via ampicillin resistance; iii/ bacterial origins of replication that produce high copy number and high plasmid yield upon extraction from bacteria; iv/ a large polylinker or multiple cloning site (MCS) with a selection of unique endonuclease cut sites to enable insertion of new DNA sequences; v/ T3 and T7 phage promoters flanking the MCS to permit *in vitro* RNA transcription; and vi/ a β-galactosidase coding region overlapping the MCS to permit colorimetric screening of bacterial colonies for successful integration of DNA insert.

**Figure 1.**
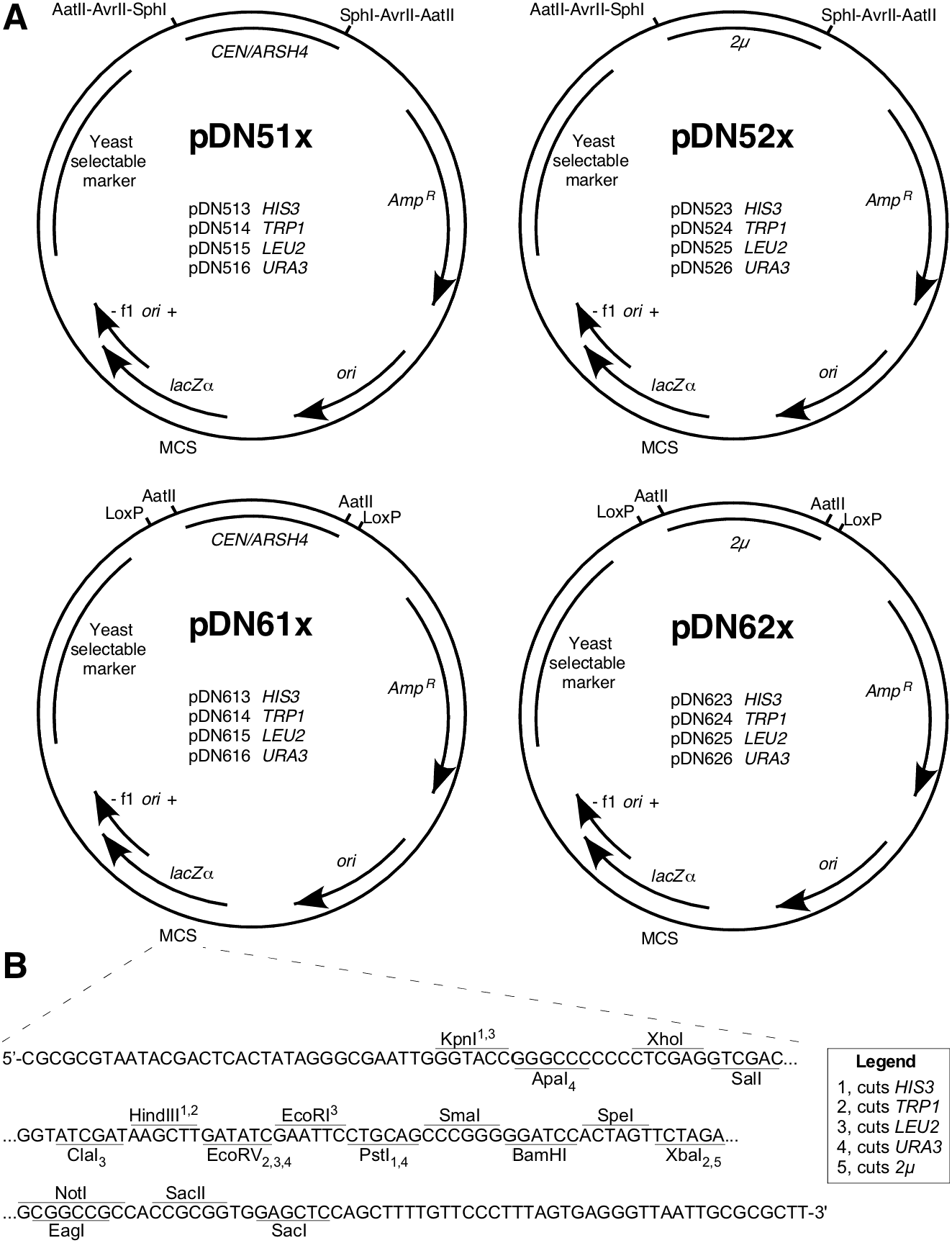
Functional maps for pDN5xx and pDN6xx series of low-copy and high-copy vectors. A) Maps displaying consistent architectural features and specific functional differences of pDN5xx and pDN6xx families. Selected restriction enzyme cut sites and LoxP sequences flanking replication loci are displayed. B) Multiple cloning site (MCS) in focus, displaying nucleotide sequence of single strand (template strand for *lacZα*, β-galactosidase) and unique restriction enzyme cut sites. Subscript and superscript numerals with each enzyme indicate capacity for enzyme to cut yeast selectable marker loci as indicated in legend. *CEN*, centromeric sequence, low copy yeast replication locus. *ARSH4*, autonomous replicating sequence. 2μ, yeast high copy replication locus. *Amp*^*R*^, ampicillin resistance gene (β-lactamase). *ori*, high copy *E. coli* origin or replication. f1 *ori*, f1 bacteriophage origin of replication. Plasmid loci depicted at approximate scale. Full plasmid sequences and annotated maps are available in Supplementary Materials.

In designing the pDN500 and −600 series (Figure 1 A), we honored the original numbering convention of the pRS series in which the second numeral indicates the yeast replication locus (‘0’ for none, ‘1’ for centromeric, and ‘2’ for 2μ) and the third indicates the yeast selectable marker (‘3’ for *HIS3*, ‘4’ for *TRP1*, ‘5’ for *LEU2*, and ‘6’ for *URA3*). In the pDN500-series, yeast replication loci are flanked by pairs of restriction endonuclease cut sites, enabling targeted *in vitro* removal of either *CEN* or *2μ* loci. The trio of AatII, AatII and SphI were selected because these enzymes are commonly used for laboratory cloning and lack cut sites elsewhere in the pRS and pDN family of vectors, excepting that AvrII cuts in the *HIS3* locus, consistent with characterization of the pRS vectors by (Chee and Haase 2012). Three pairs of flanking endonuclease cut sites were included to maximize the likelihood that at least one pair of cut sites should remain available to modify the replication locus in case further DNA inserts might contain cut sites for one or two of these enzymes. Importantly, removal of the replication locus by flanking restriction digest supports either elimination of the locus by plasmid re-circularization or substitution of an alternative replication locus.

The pDN61x and -62x series of vectors allow remodeling of yeast replication loci via a pair of parallel, flanking LoxP sequences, offering options to remodel the plasmid by Cre recombinase activity either *in vivo* or *in vitro.* The Cre-Lox recombination strategy offers the further advantage that the option to remodel a yeast replication locus will persist regardless of what further DNA sequences might be inserted into the plasmid, provided no additional LoxP sites are included. pDN600-series vectors also include a pair of flanking AatII cut sites on either side of the replication loci, so remodeling options available to the pDN500-series also apply.

### Conversion of replicating plasmids to integrating plasmids using Cre recombinase

We demonstrated working procedures for remodeling replication loci in the pDN600-series using plasmid pJM1 (Figure 2 A), generated by incorporating a gene (*GFP-CPS*) encoding GFP-tagged transmembrane endosomal cargo reporter carboxypeptidase S into the centromeric, uracil-selected vector pDN616. Chromosomal integration is particularly useful for plasmids encoding fluorescent reporters, since uneven expression across a population of cells can result in a range of signal intensities and possible phenotypes. We converted pJM1 into a replication-deficient, integrating plasmid (pJM3) via Cre recombinase activity toward the LoxP-flanked *CEN/ARSH4* locus. We passaged pJM1 through *E. coli* strain N2114Sm (Seifert *et al.* 1986) that stably expresses Cre. Plasmid DNA was extracted from ampicillin-selected N2114Sm colonies and used to transform *E. coli* strain TOP10F’ for the dual purpose of filtering the polyclonal plasmid population immediately derived from N2114Sm and achieving a higher yield of plasmid DNA. Restriction analysis of the resulting pJM3 candidates revealed that neither candidate retained the AatII cut sites that flank the *CEN/ARSH4* locus in pJM1 (Figure 2 B). Digestion with a combination of EcoRV and PvuI further confirmed that a pJM1 restriction product containing the *CEN/ARSH4* locus (1834 bp) was absent in pJM3 candidates, but that the restriction fragment at ~1250 bp in pJM3 had doubled in relative intensity, indicating the presence of two fragments, consistent with the 1834 bp fragment being reduced to 1250 bp by removal of the *CEN/ARSH4* sequence by successful Cre-Lox recombination.

**Figure 2.**
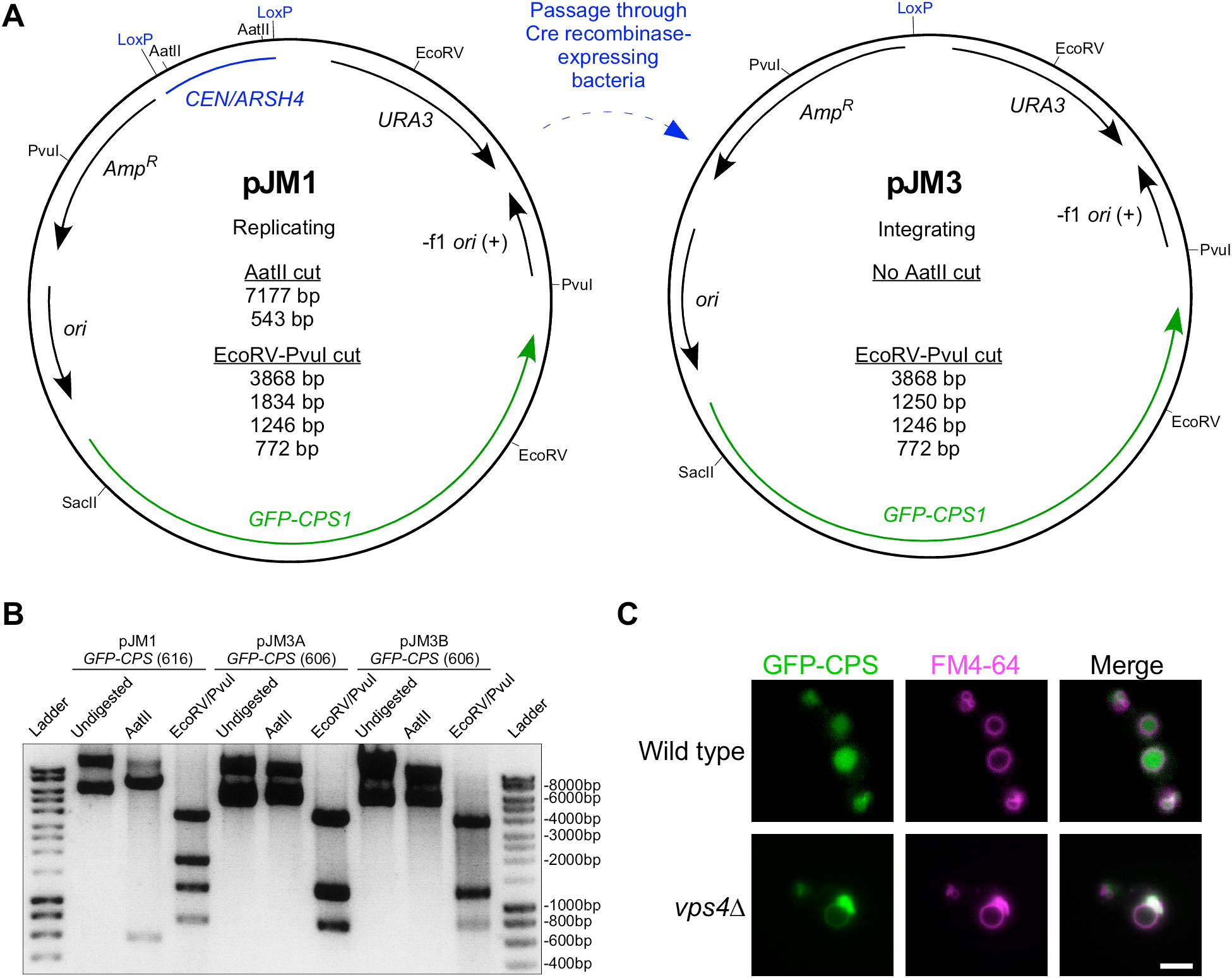
Conversion of replicating, episomal vector to integrating vector via Cre recombinase. A) Plasmid maps of low copy replicating (pJM1) and integrating (pJM3) plasmids, including relevant endonuclease enzyme cut sites and predicted restriction fragment product sizes. B) Agarose gel electrophoretic analysis of restriction digest products derived from pre-Cre-treated plasmid pJM1 and post-Cre-treated plasmid pJM3 candidates A and B. Note that 1246 bp and 1250 bp fragments predicted from EcoRV-PvuI double digests of pJM3 appear as a single band. Undigested and uncut plasmids show high molecular weight bands representing supercoiled, nicked, and concatenated circular DNA whose gel migration should not be compared to linear ladder size standards. Unlabeled DNA ladder bands are of length halfway between neighboring labeled bands. C) Fluorescence microscopy of FM 4-64-labeled, logarithmic phase yeast expressing chromosomally integrated pJM3. pJM3 was digested with SacII to integrate in the chromosomal *PCR1* promoter. Chromosomal integrants were selected on media lacking uracil. Cells were cultured in non-selective media prior to imaging. Scale bar = 1 μm.

We confirmed competence of pJM3 as an integrating plasmid by cutting at the unique SacII recognition site in the *PRC1* promoter sequence driving expression of *GFP-CPS*, producing linearized pJM3 with end sequences to direct integration via homologous recombination into the chromosomal *PRC1* promoter. We transformed linearized pJM3 into wild type yeast and a *vps4Δ* mutant strain that suffers an endosome maturation defect preventing formation of luminal vesicles (Babst *et al.* 1997), confirmed integration in colonies selected on media lacking uracil, and cultured cells under non-selective conditions prior to pulse-labeling with a fluorescent endocytic membrane dye, FM4-64 (Vida and Emr 1995) to stain the perimeter of the yeast vacuole. GFP-CPS transits via the biosynthetic pathway from Golgi to endosome where it is sorted into luminal endosomal vesicles (Odorizzi *et al.* 1998). Luminal vesicles and GFP-CPS cargo are delivered to the vacuole lumen when endosomes fuse with the vacuole, so in wild type cells GFP signal appeared inside the FM4-64-stained vacuole membrane (Figure 2 C). Loss of the gene *VPS4* (*vps4Δ*) disrupts the ability of endosomes to invaginate and form luminal vesicles, so GFP-CPS remains at the outer membrane of endosomes and is delivered instead to the outer membrane of the vacuole, co-localizing with FM4-64 at the vacuole membrane and at perivacuolar endosomal compartments (Figure 2 C). These observations are consistent with previous studies (Odorizzi *et al.* 1998) and confirm performance of the pDN600 series in converting to integration-competent vectors.

We further examined whether Cre would remove LoxP-flanked 2μ replication loci. We passaged the high copy vector pDN624 through Cre-expressing bacteria via the same procedure described above. Restriction analysis using endonuclease EcoRI revealed that all Cre-treated candidates had been reduced in length ~1400 bp compared to untreated pDN624 (Figure 3 A), consistent with the predicted loss of 1390 bp due to Cre recombination of the *LoxP::2μ::LoxP* cassette. Purified Cre enzyme is readily available from commercial suppliers, so we also examined whether LoxP-flanked yeast replication loci could be removed *in vitro*. Cre-treatment of high copy plasmids pDN624 and pDN626 for only thirty minutes resulted in modification of a substantial subpopulation of the plasmids (Figure 3 B). Treated samples possessed bands representing unmodified pDN624 and pDN626 as well as an additional band ~1400 bp shorter in length, consistent with the predicted loss of 1390 bp from each after *LoxP::2μ::LoxP* recombination. Users of the pDN600 series therefore have the option to conduct Cre-mediated remodeling of the yeast replication locus either *in vivo* or *in vitro*.

**Figure 3.**
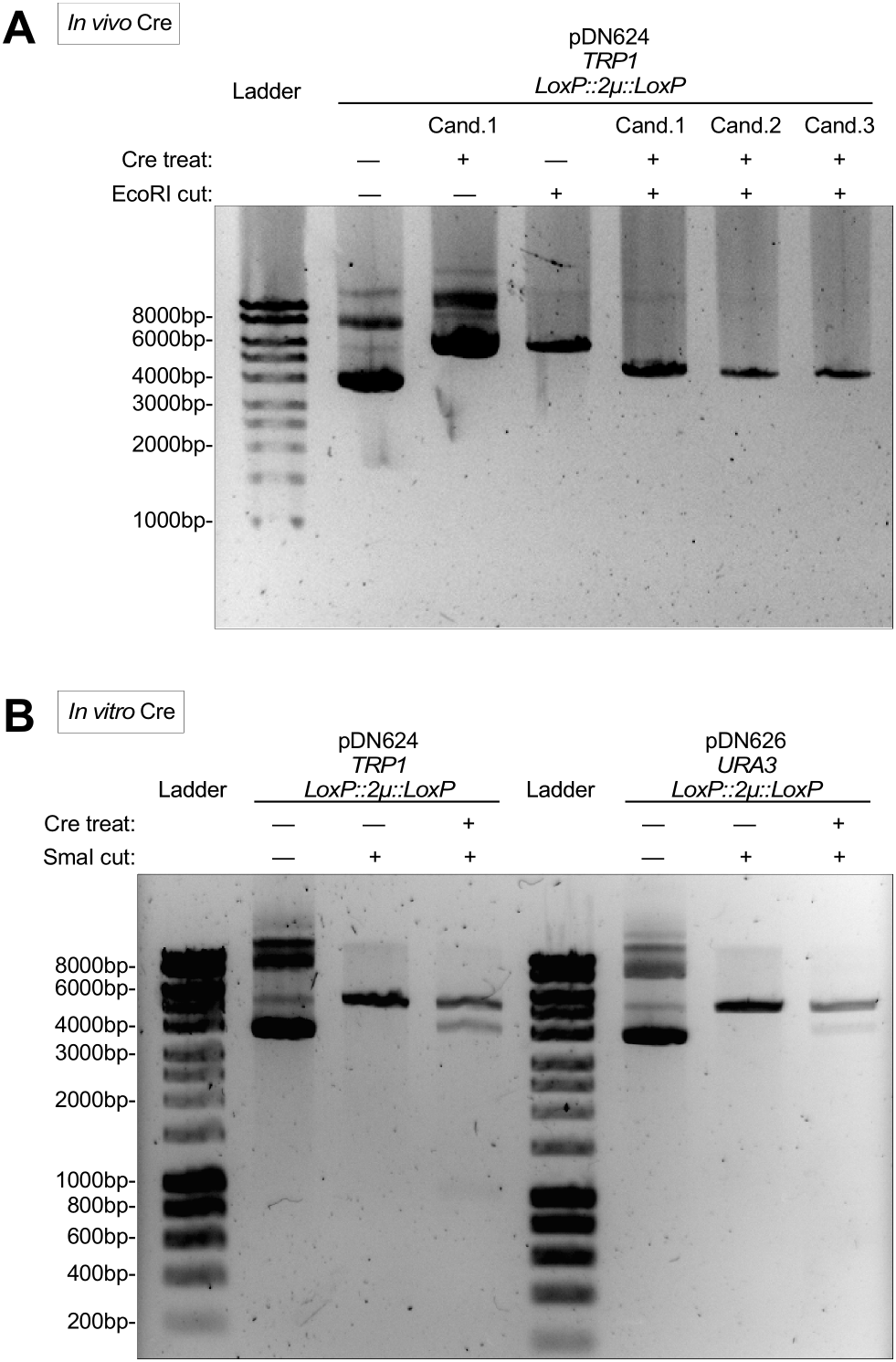
Cre-mediated removal of *2μ* replication locus *in vivo* and *in vitro*. A) Agarose gel electrophoretic confirmation of removal of *2μ* replication locus after passage of pDN624 through Cre-expressing bacterial strain N2114Sm. Each candidate represents a unique plasmid isolate from a single N2114Sm pDN624 colony. EcoRI-cut (linearized) pDN624 produces predicted bands of 5692 bp and 4302 bp before and after removal of *2μ* locus, respectively. B) Agarose gel electrophoretic confirmation of removal of *2μ* replication loci after *in vitro* treatment of pDN624 and pDN626 with Cre recombinase. Cre-treated samples represent polyclonal populations that include both unmodified (5692 bp for pDN624; 5802 bp for pDN626) and modified plasmids (4302 for pDN624; 4412 for pDN626). Unlabeled DNA ladder bands are of length halfway between neighboring labeled bands.

### Generation of integrating plasmids by restriction digest to excise plasmid replication loci

We explored the plasticity of the pDN500-series by generating a full suite of replicating and integrating plasmid vectors expressing a mutant allele (*L291A, L292A—or ‘LALA’*) of the SNARE disassembly adaptor protein Sec17 (Schwartz *et al.* 2017). We generated a *sec17*^*LALA*^ PCR product with ends homologous to the ends of SacI-digested pDN516, inserting *sec17*^*LALA*^ into pDN516 via co-transformation into yeast cells for plasmid gap repair by homologous recombination (Figure 4 A). The resulting low copy, centromeric plasmid (pDN366) was subsequently converted to a non-replicating, integrating plasmid by digesting pDN366 with AatII and re-circularizing with T4 ligase to remove the *CEN/ARSH4* locus, resulting in pDN370. In order to generate a high copy plasmid to overexpress *sec17*^*LALA*^, pDN370 was again cut with AatII to make linear ends available for insertion of the *2μ* locus. We examined efficiency of *2μ* locus insertion into pDN370 via Gibson assembly (Gibson *et al.* 2009), which like gap repair cloning in yeast also relies upon overlapping homologous sequences at the ends of vector and insert, resulting in pDN369.

**Figure 4.**
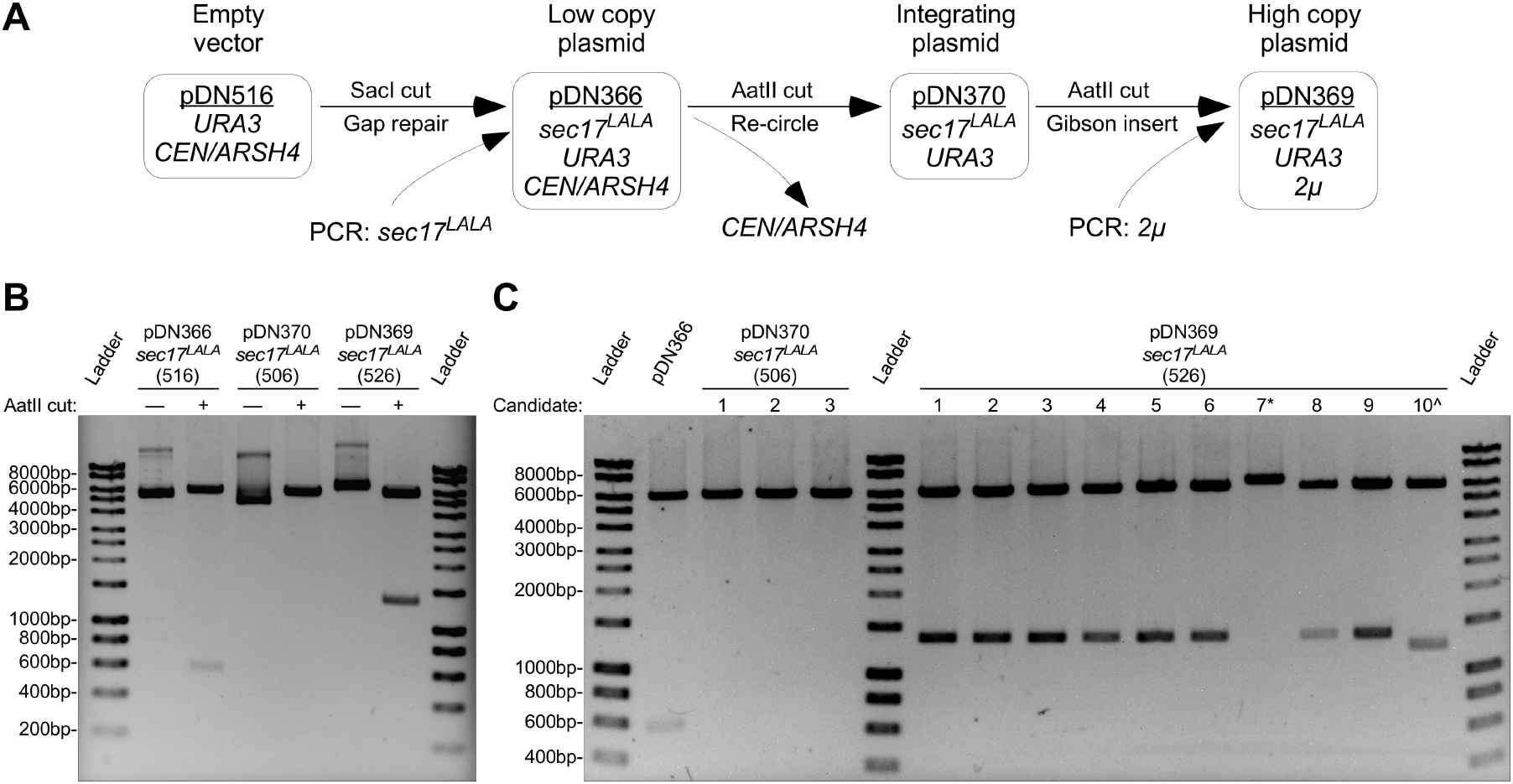
Example workflow to generate low copy, high copy, and integrating plasmids from a common precursor. A) Workflow schematic representing modification of original low copy replicating vector by insertion of a PCR product at MCS, followed by removal of original replication locus and replacement with high copy replication locus. B) Agarose gel electrophoresis and restriction enzyme analysis of plasmids resulting from demonstrated workflow. Observed AatII restriction fragments conform to predicted sizes: 6185 bp and 563 bp for pDN366; 6185 bp for pDN370; and 6185 bp and 1369 bp for pDN369. C) Agarose gel electrophoretic analysis of efficiency of removal of *CEN/ARSH4* replication locus and replacement with *2μ* replication locus. All samples shown were digested with AatII to linearize vector (no replication locus) or cut on either side of replication locus. * and ^ symbols indicate failed pDN369 candidates. Unlabeled DNA ladder bands are of length halfway between neighboring labeled bands.

All three versions of the *sec17*^*LALA*^ plasmid suite produce a 6185 bp band upon AatII digestion (Figure 4 B), representing the *sec17*^*LALA*^ gene insert and the remainder of the common plasmid backbone that lacks the yeast replication locus. pDN366 and pDN369 also produce AatII restriction fragments at 563 or 1369 bp, representing *CEN/ARSH4* or *2μ* loci, respectively. Candidate plasmids examined after recircularization of pDN366 to produce pDN370 all lack the 563 bp *CEN/ARSH4* locus (Figure 4 C), which was expected given the known robustness of the recircularization technique. Screening ten candidate plasmids for high copy pDN369 generated by Gibson assembly also revealed a high degree of successful *2μ* locus insertion (80%).

The workflow described in Figure 4 works equivalently if the starting vector is high copy (*2μ*) instead of low copy (*CEN/ARSH4*); such an alternative workflow was used to generate LUCID family cargo transport reporter plasmids (Nickerson *et al.* 2012; Nickerson and Merz 2015) in which multiple genes expressing chimeric reporter enzymes were inserted at the MCS, so the ability to remodel the replication loci of the plasmids was far preferable to subcloning large, multi-component gene inserts.

A further convenience of a standard family of yeast shuttle vectors with ability to remodel the replication loci is the limited number of needed reagents for a research lab to stock. Indeed, remodeling operations demonstrated in this study could be performed using the enzymes AatII, Cre, and T4 ligase, plus frozen stocks of *CEN/ARSH4* and *2μ* PCR products ready to insert.

Future improvements to shuttle vector systems could merge the benefits of targeted removal or remodeling of both the yeast replication loci and the yeast selectable markers. The use of Cre-Lox recombination in the pDN600-series presents an incompatibility with the commonly used Cre-Lox removal of selectable marker loci, but there are several alternative recombinase enzymes and recognition sequences available to incorporate.

## MATERIALS & METHODS

### Media and reagents

Standard methods were used for culture and storage of yeast and bacteria (Guthrie and Fink 2002). All media and reagents were purchased from Sigma-Aldrich (Saint Louis, MO) or Thermo Fisher (Waltham, MA), unless otherwise specified. All enzymes were purchased from New England Biolabs (Ipswich, MA) unless otherwise specified. High fidelity KOD Hot Start polymerase was purchased from Novagen/EMD Millipore (Darmstadt, Germany). DNA restriction digests, T4 DNA ligase reactions and PCR reactions were all performed according to manufacturer instructions. Oligonucleotide synthesis was performed by Integrated DNA Technologies (Corralville, IA). Hyperladder I (Bioline) was used as linear DNA size standard.

### DNA manipulations and reagents

Strains and plasmids used in this study are described in Table I. Oligonucleotides used in this study are described in Table II. In constructing the pDN51x or pDN52x plasmid series, PCR primer pairs (DN652p & DN653p or DN1016p & DN1017p) were designed to amplify either the *CEN/ARSH4* locus or the high copy *2μ* locus from pRS415 or pRS425 templates, respectively, while incorporating AatII, AvrII, and SphI restriction sites (‘3X’) flanking the loci. To construct the pDN61x plasmid series, PCR primer pairs (DN648p & DN649p or MQ1p & MQ2p) were designed to amplify *CEN/ARSH4* or *2μ* loci from pRS415 or pRS425 templates, respectively, while incorporating parallel LoxP sequences flanking the loci. After successful high-fidelity PCR amplification, template plasmid DNA was degraded by treatment with DpnI restriction enzyme prior to precipitation of PCR product and resuspension in 0.1M LiOAc 2mM Tris pH 7.9. The resulting *LoxP::CEN/ARSH4::LoxP*, *LoxP::2μ::LoxP, 3X::CEN/ARSH4::3X* and *3X::2μ::3X* PCR products were co-transformed into yeast (*S. cerevisiae*) with AatII-digested pRS403, pRS404, pRS405 or pRS406 linearized vectors using a lithium acetate-based protocol described below. Homologous recombination of the replication loci and linearized integrating vectors yielded new, low- and high-copy replicating vectors. pDN5xx- and pDN6xx-series plasmid candidates were screened and confirmed by restriction digest, DNA sequencing, competence for Cre-mediated recombination, and ability to support yeast and bacterial colony growth after transformation and plating onto selective media.

Yeast high-efficiency DNA transformation protocol for recircularization of linearized plasmid vectors by homologous recombination with compatible DNA insert was adapted from (Gietz *et al.* 1992). Cells were shaken overnight in YPD media at 30°C. Saturated cultures were diluted to OD_600_ ~0.1 and shaken under identical conditions until cells reached log phase density (OD_600_ = 0.4-0.6). Cells were collected by low speed centrifugation and rinsed in 0.1M LiOAc 2mM Tris pH 7.9. Cell pellets were resuspended in 50 μL 0.1M LiOAc 2mM Tris pH 7.9 containing either resuspended PCR product (‘insert’) or no PCR product as a negative control. Cell suspensions were further supplemented with 10-50 μg boiled salmon sperm DNA and approximately 25 ng linearized plasmid vector before dilution with 700 μL 40% PEG (w/v) in 0.1M LiOAc 2mM Tris pH 7.9. Cell suspensions were vortexed 10-20 seconds and incubated at 30°C for 15-30 minutes. Cell suspensions were supplemented with 5% (v/v) DMSO and vortexed another 10 seconds prior to a heat shock incubation at 42°C for 30 minutes. Cells were collected by low speed centrifugation and resuspended in 0.1M LiOAc 2mM Tris pH 7.9 prior to spreading on selective agar media.

Plasmid DNA was recovered from yeast by DNA extraction using the ‘smash and grab’ protocol (Rose *et al.* 1990) of glass bead cell lysis, phenol-chloroform extraction and ethanol precipitation of the aqueous phase to yield genomic and plasmid DNA. Plasmids were separated from genomic DNA by electroporation of *E. coli* and plating of cells to LB agar media with ampicillin (100 μg/mL). Plasmids were recovered from bacteria using a QIAgen plasmid miniprep kit (Qiagen, Valencia, CA). DNA sequencing reactions of replication loci using flanking primers DN661p or DN837p was performed by Genewiz (South Plainfield, NJ). Sequencing alignments were performed using SnapGene software (GSL Biotech, San Diego, CA).

All restriction endonuclease digests of plasmids were performed according to manufacturer’s instructions (New England Biolabs).

Cre-mediated removal of *LoxP*-flanked replication loci from pDN61x- and pDN62x-series vectors was accomplished *in vivo* by chemical transformation of an *E. coli* strain expressing Cre recombinase, N2114Sm (Seifert *et al.* 1986). Ampicillin-selected colonies were picked and grown in LB media supplemented with ampicillin (50 μg/mL) before plasmid extraction via QIAgen plasmid miniprep. Plasmid yields from N2114Sm host strain are low, so purified plasmid candidates were further passaged through TOP10F’ *E. coli* (Invitrogen, Carlsbad, CA) via chemical transformation and QIAgen plasmid miniprep extraction.

Cre-mediated removal of *LoxP*-flanked replication loci from pDN62x-series vectors was accomplished *in vitro* treating 250 ng plasmid with purified Cre recombinase enzyme (New England Biolabs) in 50 μL reaction per manufacturer instructions, incubating at 37°C for 30 minutes. Resulting polyclonal Cre recombinase reactions were heat inactivated and plasmid DNA was precipitated twice using ethanol prior to restriction digest and electrophoretic analysis.

Plasmid pJM1 was constructed via PCR amplification of the *PRC1* promoter-driven *GFP-CPS1* cassette from plasmid template pGO45 (Odorizzi *et al.* 1998) using primers DN680p and DN693p, creating a PCR product with ends overlapping the ends of PvuII-cut vector pDN616. PCR template was eliminated by DpnI digestion. PvuII cuts on either side of the multiple cloning site (MCS), removing the MCS entirely, thus eliminating many redundant endonuclease cut sites. PCR product and PvuII-cut pDN616 were co-transformed into yeast to perform homologous recombination plasmid repair as described above. Transformants were selected on agar media lacking uracil to select circularized plasmids. pJM1 was modified to generate pJM3 by removal of the LoxP-flanked *CEN/ARSH4* locus by passaging pJM1 through Cre-expressing strain N2114Sm as described above. pJM3 was linearized for chromosomal integration by SacII digestion of the unique cut site in the *PRC1* promoter sequence, generating linear ends capable of mediating homologous recombination at the *PCR1* chromosomal locus. Strains SEY6210 and MBY3 were both transformed using SacII-cut pJM3 using the high-efficiency protocol described above.

Plasmids expressing the *L291A, L292A* mutant of *SEC17* (*sec17*^*LALA*^) driven by 500 bp native *SEC17 promoter* were constructed via an overlap extension PCR scheme in which the overlapping sequences of primers DN982p and DN983p introduced the *LALA* mutation. Flanking primers DN927p and DN928p included 35 and 34 bp, respectively, sequence overlapping the linear ends of SacI-digested pDN516, allowing insertion of *sec17*^*LALA*^ PCR product into the pDN516 MCS via homologous recombination plasmid repair, resulting in pDN366. AatII digest of pDN366 linearized the plasmid and removed the *CEN/ARSH4* locus. Compatible AatII overhangs were ligated together using T4 ligase, omitting the *CEN/ARSH4* locus and resulting in pDN370. Insertion of *2μ* replication locus PCR product into AatII-digested pDN370 via Gibson cloning was performed essentially as described (Gibson *et al.* 2009), resulting in pDN369. PCR primers DN652p and DN653p used to amplify the *2μ* locus include 30 bp overhangs homologous to ends of AatII-digested pDN5xx vectors, making them suitable to mediate both homologous recombination plasmid repair in yeast and *in vitro* overlap extension via Gibson cloning.

### Imaging

Pulse-chase labeling of log phase yeast with vacuolar fluorescent dye FM4-64 and fluorescence microscopy imaging were performed as described (Nickerson *et al.* 2012), except that cells were grown in non-selective YPD prior to labeling and imaging. DNA in agarose gels was stained using ethidium bromide and visualized using ultraviolet light in a BioRad Chemi-doc system with digital camera and Quantity One imaging software (BioRad, Hercules, CA). All gel images were exported from Quantity One as .TIFF images, except Figure 2 panel B, which was printed to photographic paper and later scanned in .TIFF format using an Epson flatbed scanner. Images were cropped using Adobe Photoshop CS6 (Adobe, San Jose, CA). Fluorescence microscopy images were overlaid using ImageJ (NIH, https://imagej.nih.gov/ij/). Images were arranged as annotated figures using Canvas Draw 4 vector graphics software (Canvas GFX, Boston, MA).

## Supplementary materials and reagent requests

Vector sequence files and maps may be downloaded in SnapGene format (.dna) as supplementary files. Plasmid vectors in the pDN51x, pDN52x, pDN61x, and pDN62x vector series will be deposited at AddGene (Cambridge, MA). Direct reagent requests to the Nickerson Lab should please provide a self-addressed, stamped envelope, mailed to ‘Attn: Nickerson, 5500 University Pkwy, CSUSB Biology Dept, Rm BI-302, San Bernardino, CA 92407-2318.’

## AUTHOR CONTRIBUTIONS

DPN conceived of the project. DPN and MAQ conceived and designed experiments. DPN, JMM and MAQ performed experiments and analyzed results. DPN wrote the manuscript with editing contributions from MAQ and JMM.

## ACKNOWLEDGMENTS

This work was supported by grants from the CSUSB Office of Student Research to MAQ and DPN, charitable donations to the Nickerson Lab Research Fund (CSUSB), and NIH/NIGMS RO1 GM077349 to AJM (University of Washington).

---

**Table I.** Plasmids and strains used in this study.

| Name | Genotype/Description | Source/Reference |
| --- | --- | --- |
| *Plasmids* |  |  |
| pDN613 | Amp<sup>R</sup> *HIS3* *LoxP*::*CEN/ARSH4*::*LoxP* | This study |
| pDN614 | Amp<sup>R</sup> *TRP1* *LoxP*::*CEN/ARSH4*::*LoxP* | (Shideler *et al.* 2015) |
| pDN615 | Amp<sup>R</sup> *LEU2* *LoxP*::*CEN/ARSH4*::*LoxP* | (Paulsel *et al.* 2013) |
| pDN616 | Amp<sup>R</sup> *URA3* *LoxP*::*CEN/ARSH4*::*LoxP* | (Nickerson *et al.* 2012) |
| pDN623 | Amp<sup>R</sup> *HIS3* *LoxP*::2μ::*LoxP* | This study |
| pDN624 | Amp<sup>R</sup> *TRP1* *LoxP*::2μ::*LoxP* | This study |
| pDN625 | Amp<sup>R</sup> *LEU2* *LoxP*::2μ::*LoxP* | This study |
| pDN626 | Amp<sup>R</sup> *URA3* *LoxP*::2μ::*LoxP* | This study |
| pDN513 | Amp<sup>R</sup> *HIS3* 3X::*CEN/ARSH4*::3X | This study |
| pDN514 | Amp<sup>R</sup> *TRP1* 3X::*CEN/ARSH4*::3X | This study |
| pDN515 | Amp<sup>R</sup> *LEU2* 3X::*CEN/ARSH4*::3X | This study |
| pDN516 | Amp<sup>R</sup> *URA3* 3X::*CEN/ARSH4*::3X | This study |
| pDN523 | Amp<sup>R</sup> *HIS3* 3X::2μ::3X | This study |
| pDN524 | Amp<sup>R</sup> *TRP1* 3X::2μ::3X | (Lobingier *et al.* 2014) |
| pDN525 | Amp<sup>R</sup> *LEU2* 3X::2μ::3X | This study |
| pDN526 | Amp<sup>R</sup> *URA3* 3X::2μ::3X | (Nickerson *et al.* 2012) |
| pJM1 | Amp<sup>R</sup> *URA3* *LoxP*::*CEN/ARSH4*::*LoxP* *PRC1pr*::*GFP-CPS1* | This study |
| pJM3 | Amp<sup>R</sup> *URA3* *LoxP* *PRC1pr*::*GFP-CPS1* | This study |
| pDN314 | Amp<sup>R</sup> *URA3* 3X::2μ::3X *SEC17* | (Lobingier *et al.* 2014) |
| pDN366 | Amp<sup>R</sup> *URA3* 3X::*CEN/ARSH4*::3X *sec17*<sup>L291A/L292A</sup> | This study |
| pDN369 | Amp<sup>R</sup> *URA3* 3X::2μ::3X *sec17*<sup>L291A/L292A</sup> | (Schwartz *et al.* 2017) |
| pDN370 | Amp<sup>R</sup> *URA3* *sec17*<sup>L291A/L292A</sup> | This study |
| pRS403 | Amp<sup>R</sup> *HIS3* | (Sikorski and Hieter 1989) |
| pRS404 | Amp<sup>R</sup> *TRP1* | (Sikorski and Hieter 1989) |
| pRS405 | Amp<sup>R</sup> *LEU2* | (Sikorski and Hieter 1989) |
| pRS406 | Amp<sup>R</sup> *URA3* | (Sikorski and Hieter 1989) |
| pRS415 | Amp<sup>R</sup> *LEU2* *CEN/ARSH4* | (Sikorski and Hieter 1989) |
| pRS425 | Amp<sup>R</sup> *LEU2* 2μ | (Christianson *et al.* 1992) |
| pGO45 | Amp<sup>R</sup> *URA3* 2μ *PRC1pr*::*GFP-CPS1* | (Odorizzi *et al.* 1998) |
| *E. coli* |  |  |
| TOP10F' | *[lacI<sup>q</sup> Tn10(tet<sup>R</sup>)] mcrA $\Delta$(*mrr-hsdRMS-mcrBC*)ϕ80*lacZ* ΔM15 Δ*lacX74* *deoR* *nupG* *recA1* *araD139*Δ(*ara-leu*)7697 *galU* *galK* *rpsL*(*Str<sup>R</sup>*) *endA1* $\lambda^-$* | Invitrogen |
| N2114Sm | F<sup>-</sup> *recA* λ-*cre* *rpsL* | (Seifert *et al.* 1986) |
| *S. cerevisiae* |  |  |
| SEY6210 | *MATα* *leu2-3,112* *ura3-52* *his3-200* *trp1-901* *lys2-801* *suc2-9* | (Robinson *et al.* 1988) |
| MBY3 | SEY6210 *vps4Δ*::*TRP1* | (Babst *et al.* 1997) |
| JMY1 | SEY6210 *GFP-CPS1* (pJM3::*URA3*) | This study |
| JMY2 | MBY3 *GFP-CPS1* (pJM3::*URA3*) | This study |

**Table II**. Primer sequences used in this study. Sequences to mediate homologous recombination shown in **bold**. LoxP sequences shown in *italics*. Restriction site sequences shown as <u>underlined</u>. PCR annealing sequences shown in lowercase.

DN648p: Rev, PCR amplify *CEN/ARSH4* sequence, incorporate LoxP sequence and AatII restriction site, mediate repair of AatII-digested pRS40x vector. 5’**GGTTAATGTCATGATAATAATGGTTTCTTA***ATAACTTCGTATAGCATACATTATACGAAGTTAT*<u>GACGTC</u>ggacggatcgcttgcctg-3’

DN649p: Fwd, PCR amplify *CEN/ARSH4* sequence, incorporate LoxP sequence and AatII restriction site, mediate repair of AatII-digested pRS40x vector. 5’**CGCGCACATTTCCCCGAAAAGTGCCACCT***ATAACTTCGTATAATGTATGCTATACGAAGTTATT*<u>GACGTC</u>cccgaaaagtgccacctg-3’

DN652p: Fwd, PCR amplify *CEN/ARSH4* sequence, incorporate triplicate restriction sites, mediate repair of AatII-digested pRS40x or pDN50x vectors. 5’**CACATTTCCCCGAAAAGTGCCACCT**<u>**GACGT**CCTAGGCATGC</u>ggtccttttcatcacgtgc-3’

DN653p: Rev, PCR amplify *CEN/ARSH4* sequence, incorporate triplicate restriction sites, mediate repair of AatII-digested pRS40x or pDN50x vectors. 5’**ATGTCATGATAATAATGGTTTCTTA**<u>**GACGT**CCTAGGCATGC</u>gataataatggtttcttag-3’

DN1016p: Fwd, PCR amplify *2μ* sequence, incorporate triplicate restriction sites, mediate repair of AatII-digested pRS40x or pDN50x vectors. 5’**CACATTTCCCCGAAAAGTGCCACCT**<u>**GACGT**CCTAGGCATGC</u>aacgaagcatctgtgcttcatt-3’

DN1017p: Rev, PCR amplify *2μ* sequence, incorporate triplicate restriction sites, mediate repair of AatII-digested pRS40x or pDN50x vectors. 5’**ATGTCATGATAATAATGGTTTCTTAGACGT**CCTAGGCATGCgatccaatatcaaaggaaatg-3’

MQ1p: Fwd, PCR amplify *2μ* locus, mediate insertion into AatII-digested pDN61x family vectors to create pDN62x family vectors. 5’**ATAATGTATGCTATACGAAGTTATT**<u>**GACGT**c</u>aacgaagcatctgtgcttcattttg-3’

MQ2p: Rev, PCR amplify *2μ* locus, mediate insertion into AatII-digested pDN61x family vectors to create pDN62x family vectors. 5’**TATAGCATACATTATACGAAGTTAT**<u>**GACGT**c</u>gatccaatatcaaaggaaatgatagc-3’

DN661p: Sequencing primer for yeast replication loci, anneals near yeast selectable marker locus. 5’tacaatctgctctgatgcc-3’

DN837p: Sequencing primer for yeast replication loci, anneals in *AmpR* promoter. 5’ttattgaagcatttatcaggg-3’

DN680p: Rev, PCR amplify 200 bases of *CPS1* terminator to copy *CPYpr::GFP-CPS1* cassette, mediate repair of PvuII-digested pDN616. 5’**GATCGGTGCGGGCCTCTTCGCTATTACGCCAG**taaattttgatttgacacttg-3’

DN693p: Fwd, PCR amplify 455 bases of *PRC1* promoter to copy *PRC1pr::GFP-CPS1* cassette, mediate repair of PvuII-digested pDN616. 5’**CCTCTCCCCGCGCGTTGGCCGATTCATTAATGCAG**attgacagagcagtatgtgagg-3’

DN927p: Fwd, PCR amplify 500 bp *SEC17* promoter, mediate gap repair into SacI-digested pDN516. 5’**CTAGTTCTAGAGCGGCCGCCACCGCGGTG<u>GAGCTC</u>**ttctttgtcaattgcatctcta-3’

DN928p: Rev, PCR amplify 300 bp of *SEC17* terminator, mediate gap repair into SacI-digested pDN5xx. 5’**TAACCCTCACTAAAGGGAACAAAAGCTG<u>GAGCTC</u>**ggaagatccttacattacacg-3’

DN982p: Fwd, introduce *L291A L292A* (‘LALA’) mutation into *SEC17* via sequence overlap extension PCR, overlaps with DN983p. 5’ATCCAGCAACAAGAAGATGAT GCG GCA TGA acggcatatacttacgcgca-3’

DN983p: Rev, introduce *L291A L292A* (‘LALA’) mutation into *SEC17* via sequence overlap extension PCR, overlaps with DN982p. 5’TGCGCGTAAGTATATGCCGT TCA TGC CGC atcatcttcttgttgctggat-3’

